# Convergent reductive evolution of cyanobacteria in symbiosis with Dinophysiales dinoflagellates

**DOI:** 10.1101/2024.01.11.574452

**Authors:** Takuro Nakayama, Mami Nomura, Akinori Yabuki, Kogiku Shiba, Kazuo Inaba, Yuji Inagaki

## Abstract

The diversity of marine cyanobacteria has been extensively studied due to their vital roles in ocean primary production. However, little is understood about the diversity of cyanobacterial species involved in symbiotic relationships. In this study, we successfully sequenced the complete genome of a cyanobacterium in symbiosis with *Citharistes regius*, a dinoflagellate species thriving in the open ocean. A phylogenomic analysis revealed that the cyanobacterium (CregCyn) belongs to the marine picocyanobacterial lineage, akin to another cyanobacterial symbiont (OmCyn) of a different dinoflagellate closely related to *Citharistes*. Nevertheless, these two symbionts are distinct lineages, suggesting independent origins of their symbiotic lifestyles. Despite the distinct origins, the genome analyses of CregCyn revealed shared characteristics with OmCyn, including an obligate symbiotic relationship with the host dinoflagellates and a degree of genome reduction. In contrast, a detailed analysis of genome subregions unveiled that the CregCyn genome carries genomic islands that are not found in the OmCyn genome. The presence of the genomic islands implies that exogenous genes have been integrated into the CregCyn genome at some point in its evolution. This study contributes to our understanding of the complex history of the symbiosis between dinoflagellates and cyanobacteria, as well as the genomic diversity of marine picocyanobacteria.

## Introduction

Cyanobacteria are essential primary producers that underpin the global ecosystem. Within this group, marine picocyanobacteria composed of *Prochlorococcus* and *Synechococcus* are known to contribute a significant part of the ocean net primary production (Flombaum et al., 2013), particularly in nutrient-poor open-ocean regions. For heterotrophic microorganisms inhabiting these nutrient-limited areas, the acquisition of organic matters is vital. Therefore, interactions with primary producers, such as the symbiosis with a cyanobacterium, are of great importance for survival in such harsh environments. While the diversity of marine cyanobacteria has been extensively studied in the context of marine microbial ecology, previous analyses have mainly focused on free-living cyanobacteria, leaving our understanding of species in symbiosis with other organisms limited. In recent years, previously unrecognized lineages of marine cyanobacteria have been discovered within symbiotic relationships with eukaryotes (Nakayama et al., 2019; Schvarcz et al., 2022; Thompson et al., 2012). Such findings suggest the need for further investigation of cyanobacteria that engage in symbiosis to fully understand the diversity of marine cyanobacteria.

Dinophysiales dinoflagellates are known to be among the organisms that are in a symbiotic relationship with cyanobacteria in the open ocean. The majority of the members of Dinophysiales lack plastids, except for the genus *Dinophysis*, of which members utilize plastids of other photosynthetic organisms ingested as prey (Park et al., 2014). In this order, genera such as *Ornithocercus, Histioneis, Parahistioneis, Amphisolenia*, and *Citharistes* are found in subtropical and tropical waters and have long been known to form symbiotic relationships with cyanobacteria (Lucas, 1991). Some of these symbiotic species have developed cell coverings, also called thacal plates, resulting in conspicuous cell morphologies. In addition, in the cells of *Ornithocercus, Histioneis, Parahistioneis*, and *Citharistes*, some of the thacal plates form a chamber-like space in which symbiotic cyanobacteria live. Inside this symbiotic space, which is also known as a phaeosome chamber, is topologically extracellular, and thus the cyanobacteria living in the chamber are extracellular symbionts. The existence of this special structure implies that the Dinophysiales species living in these oligotrophic oceans have evolved based on symbiosis with cyanobacteria. A recent genome analysis of symbiotic cyanobacteria living within the chamber of *Ornithocercus magnificus* has revealed its evolutionary background and metabolic characteristics (Nakayama et al., 2019). The study revealed that the symbiotic cyanobacteria of *O. magnificus* (hereafter referred to as OmCyn) belong to the marine *Synechococcus* clade and occupied the basal position of the subcluster 5.1, a previously unrecognized lineage of *Synechococcus* that, together with *Prochlorococcus*, is dominant in the ocean. A further analysis using available metagenomic data as a reference revealed that the symbiotic relationship between this cyanobacterium and the dinoflagellate host is obligate, suggesting that this cyanobacterium cannot survive alone in the ocean but is passed from chamber to chamber across host generations (Nakayama et al., 2019). However, the fact that OmCyn was vertically inherited does not imply that Dinophysiales dinoflagellates other than *Ornithocercus* have cyanobacteria of the same lineage as OmCyn. Lucas (1991) made pioneering microstructural observations of symbiotic cyanobacteria of Dinophysiales species, and classified the symbionts found in several dinoflagellate species into four types based on differences in morphology. The morphological diversity of symbiotic cyanobacteria was also reported by Foster et al. (2006a), and it was observed that in some Dinophysiales genera several morphological types of cyanobacteria were mixed within their chamber (Foster et al., 2006a). The presence of these morphological differences suggests that there is phylogenetic diversity in symbiotic cyanobacteria throughout the Dinophysiales dinoflagellates and this idea has been supported by phylogenetic analyses based on partial sequences of the 16S rDNA of these cyanobacterial symbionts (Foster et al., 2006b; Kim et al., 2021). However, because that whole genome information is available only for OmCyn, the diversity of cyanobacterial symbionts of Dinophysiales dinoflagellates, beyond OmCyn, including their detailed phylogenetic positions among free-living relatives and their genomic features, remains unclear. Such information will help us to capture the entire evolutionary history of the symbiotic relationship between cyanobacteria and Dinophysiales dinoflagellates.

*Citharistes*, another Dinophysiales dinoflagellate genus with symbiotic cyanobacteria, has been shown by molecular phylogenetic analysis to be closely related to species of the genus *Ornithocercus* (Jensen & Daugbjerg, 2009). Despite the close phylogenetic relationship, the morphology of a *Citharistes* cell has features that differ significantly from those of *Ornithocercus*. In *Ornithocercus*, the symbiotic chamber is formed by a well-developed cingular list, a structure of the thecal plate that surrounds the periphery of a cell, giving a crown-like shape (Supplementary figure S1). Although the chamber is surrounded by two cingular lists, the space is not closed, and there is enough opening for a cell with the size of a cyanobacterium to easily move in and out between the chamber and the outside environment. On the other hand, the symbiotic chamber of *Citharistes* is more closed: the chamber of *Citharistes* is developed to be embedded in the cell (Supplementary figure S1) and is connected to the external environment by a single small hole through which a cyanobacterial cell would barely pass. Such a semi-closed chamber has not been seen in other Dinophysiales dinoflagellates. Furthermore, the symbiotic cyanobacteria of *Ornithocercus* are speculated to be occasionally preyed upon by their hosts (Lucas, 1991; Nakayama et al., 2019; Tarangkoon et al., 2010), but the symbiotic chamber of *Citharistes* where the symbionts reside is spatially separated from the location of the cell mouth (near the flagellum base), which suggests that the symbiotic relationship with the host may also differ between cyanobacterial symbionts between these two dinoflagellate genera. The phylogenetic and metabolic characteristics of symbiotic cyanobacteria of *Citharistes* are not clear. Foster et al. (2006b), in a challenging attempt to directly reverse transcribe 16S rRNA from single *Citharistes* cells isolated from field samples, identified several distinct cyanobacterial sequences. However, due to the short length of the partial 16S rRNA sequences analyzed (286–379 bp) and the limited number of publicly available cyanobacterial sequences for comparison at the time of the study, little hint for the identities of their cyanobacterial symbiont could be obtained by the phylogenetic analysis. In this study, we have succeeded in completely sequencing the genome of the symbiotic cyanobacterium of *Citharistes regius* (hereafter referred to as CregCyn). CregCyn was shown to belong to the picocyanobacterial lineage like the previously known symbiotic cyanobacterium of *Ornithocercus magnificus* but is of clearly a distinct lineage. The results also suggest that the symbiosis between CregCyn and the host dinoflagellate is obligatory and that the symbiont genome has experienced genome reduction along with the intimate symbiotic relationship. These results suggest a complex evolutionary history of cyanobacterial symbiosis in Dinophysiales dinoflagellates and reconfirm the previously overlooked diversity in cyanobacteria.

## Results and Discussion

### Genome sequence of cyanobacteria isolated from a *Citharistes regius* cell

Three individuals of *Citharistes regius* (Supplementary figure S2) were found in surface seawater off Shimoda, Shizuoka, Japan, and whole genome amplification was performed for each of them. Hybrid assembly of Illumina short reads and Oxford Nanopore long reads for one of the cells revealed three contigs and a small circular sequence (∼17 Kbp), that are identified as cyanobacterial in origin. PCR-based gap-closing confirmed that the three contigs constituted a single circular chromosome of 1.94 Mbp. All the cyanobacterial sequences above were consistently detected in the sequence data of the two other *Citharistes regius* cells, leading to the conclusion that the genome originates from the symbiotic cyanobacterium of *C. regius* (designated as CregCyn; (Figure 1)). A small circular sequence of approximately 17 Kbp was thought to be a plasmid associated with this symbiotic cyanobacterium. Preliminary phylogenetic analysis of the 16S rRNA gene (Supplementary figure S3) showed that the 16S rRNA gene sequence from the CregCyn genome assembly formed a monophyletic clade with some of the partial rRNA gene sequences obtained from the *Citharistes* cells in a previous study (Foster et al., 2006b).

**Figure 1.**
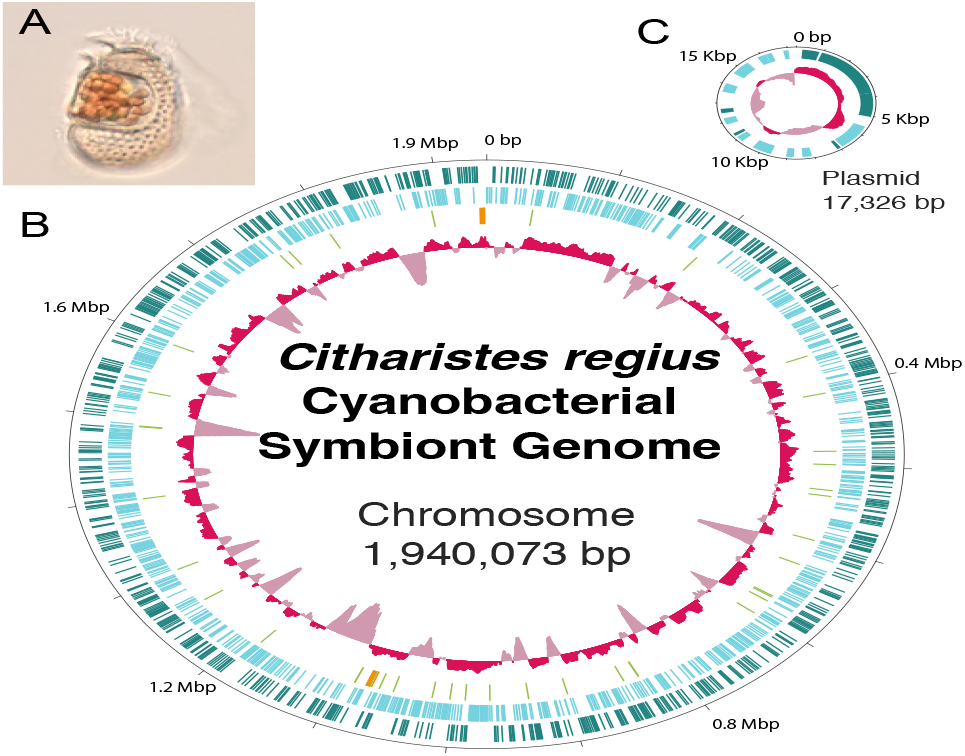
Genome map for the cyanobacterial symbiont of *Citharistes regius*. (**A**) Micrograph of *C. regius* found at the same sampling site as the individuals used in this study. Orange granular structures are cells of symbiotic cyanobacteria. (**B**) Map of the circular chromosome. The two outer circles (green and light blue) show the positions of protein-coding genes on plus and minus strands. Light green and orange bars in the third circle indicate tRNA and rRNA genes, respectively. The innermost circle shows the relative G+C content. (**C**) Map of the plasmid. Outer circle show protein-coding genes (green and light blue for plus and minus strand, respectively). The inner circle shows the relative G+C content.

### Phylogenetic relationship with another cyanobacterial symbiont of a dinoflagellate

A multigene phylogenetic analysis was conducted to investigate the phylogenetic relationship of CregCyn with OmCyn, a symbiont of *Ornithocercus magnificus*, whose phylogenetic position had already been revealed (Nakayama et al., 2019). The maximum likelihood phylogenetic tree (Figure 2) inferred from 143 protein amino acid sequences indicated that OmCyn occupied a unique phylogenetic position at the base of *Synechococcus* subcluster 5.1 (Ahlgren & Rocap, 2012), consistent with a previous study (Nakayama et al., 2019). The branch of CregCyn, on the other hand, was nested within the *Synechococcus* subcluster 5.1 and displayed a sister relationship with a monophyletic lineage named clade IV (Ahlgren & Rocap, 2012), comprising *Synechococcus* sp. CC9902 and BL107. The backbone topology for *Synechococcus* subcluster 5.1, including the two symbiotic cyanobacteria, was supported by high bootstrap values (100% or 99%) for all nodes. A previous study suggested that the hosts of the two symbionts, *Ornithocercus* and *Citharistes*, are phylogenetically closely related in Dinophysiales (Jensen & Daugbjerg, 2009). However, our phylogenomic analysis indicates that their symbionts, OmCyn and CregCyn, originate from closely related yet distinct cyanobacterial lineages.

**Figure 2.**
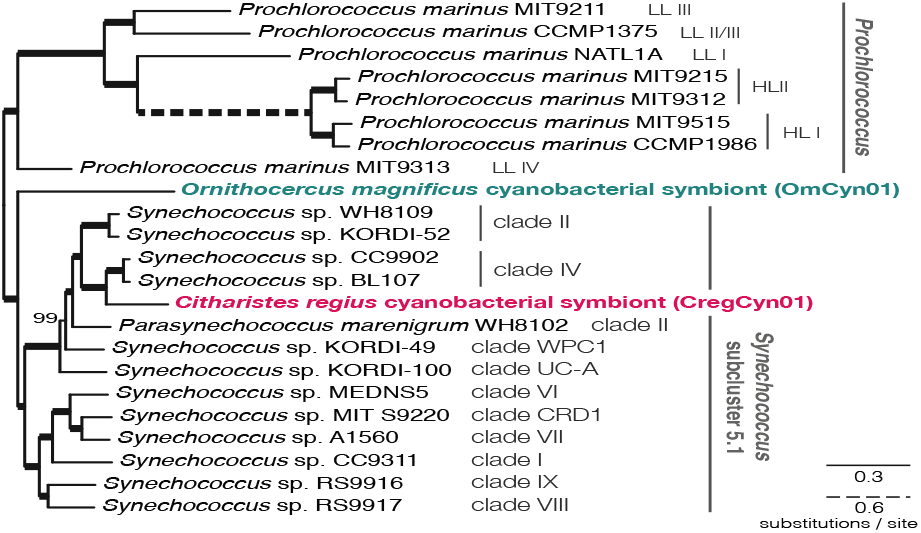
Maximum likelihood phylogenetic tree showing the positions of two cyanobacterial symbionts of dinoflagellates. A maximum likelihood phylogenetic tree inference was performed using a concatenated protein alignment of 143 protein sequences from 23 species/strains, under the site-heterogeneous substitution model LG+C20+F+Γ4. Support value for each node obtained from 100 nonparametric bootstrap replicates under the LG+C20+F+Γ4-PMSF model. Thick branches indicate 100% bootstrap values. Scale bars represents the estimated number of amino acid substitutions per site

### Association with host dinoflagellates in the natural environment

To gain insight into the interaction of CregCyn with its host in the natural environment, we analyzed the occurrence pattern of the CregCyn genome in the global marine metagenome data provided by the *Tara* Oceans project (Pesant et al., 2015; Sunagawa et al., 2015). We checked the occurrence of the genomic sequences of CregCyn as well as those of OmCyn and free-living picocyanobacteria in the metagenomes of continuously size fractionated seawater samples (0.8-5 μm, 5-20 μm, 20-180 μm and 180-2,000 μm) obtained from 66 *Tara* Oceans sampling stations in a wide range of oceans (Sunagawa et al., 2015). The analysis showed that most metagenomic reads aligned with the free-living picocyanobacterial genome were found in the smallest size fraction (0.8-5 μm; Figure 3). This result is reasonable given that the cell size of picocyanobacteria is less than 2 μm in diameter. In contrast, the genomes of the cyanobacteria symbiotic with the two dinoflagellates yielded few corresponding reads from this small size fraction. In agreement with a previous study (Nakayama et al., 2019), most of the sequences homologous to the OmCyn genome were obtained from the 20-180 μm size fraction (88% of the total; Figure 3). Likewise, sequence reads aligning with the CregCyn main chromosome were predominantly detected in the 20-180 μm size fraction (63%), followed by the 5-20 μm size fraction (36%). Although the number of metagenomic reads corresponding to the plasmid sequence of CregCyn were scarce (15 out of ∼1.77 billion searched reads), all the reads were exclusively detected from the 20-180 μm size fraction. Given that the size range of 5-180 μm corresponds to eukaryotic microorganisms such as dinoflagellates, as previously discussed (Nakayama et al., 2019), the observed distribution pattern implies a tight physical association between these symbiotic cyanobacteria and their respective host cells in natural environments. This suggests that CregCyn, like OmCyn, does not live freely in the environment but rather undergoes vertical inheritance across host generations. The fact that reads corresponding to the OmCyn genome were concentrated in the 20-180 μm size fraction, whereas reads for the CregCyn genome were also detected in the 5-20 μm size fraction, may be due to the size of the host dinoflagellate. The cell size of *Ornithocercus magnificus* is approximately 75-115 μm (Okolodkov, 2014), while the cell size of *Citharistes regius* is rather small; the cell size of the individuals examined in this study is 40-50 μm even at the longest axis of the cell outline, including the ornaments formed by thacal plates (Supplementary figure S2).

**Figure 3.**
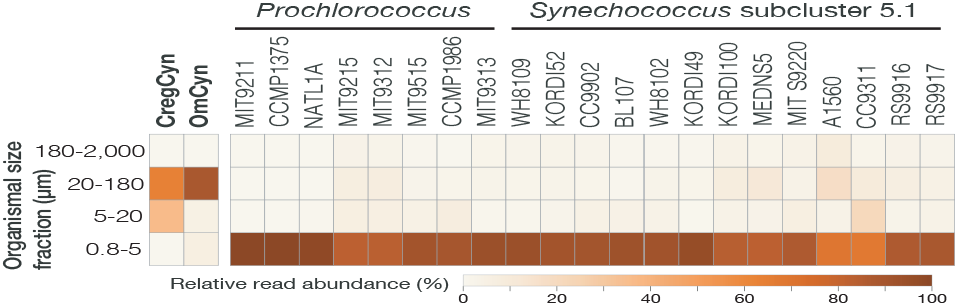
Relative metagenomic read abundances among four size fractions. A total of 570 metagenomic samples from 66 sampling sites provided by the *Tara* Oceans project were used for this estimation. Reads corresponding to the free-living cyanobacterial genomes were most abundant in the bacterialsized fractions, whereas those corresponding to the dinoflagellate symbiotic cyanobacterial genomes were obtained exclusively from the larger size fractions.

### Genome reduction in the CregCyn genome

While our phylogenetic analysis suggested phylogenetic independence, we observed a similar degree of genome reduction in CregCyn and OmCyn. While previous studies have shown that OmCyn has a reduced genome relative to the free-living picocyanobacteria genome (Nakayama et al., 2019), our comparative genomic analysis revealed that CregCyn has a compact protein repertoire comparable to that of OmCyn (Figure 4). The protein repertoires of the other free-living strains in the *Synechococcus* subcluster 5.1 range from 2,000 to 2,400, while those of the two symbionts are just over 1,500, which is only 1/4 to 1/3 the size of the closely related free-living strains. This protein repertoire is even smaller than that of *Prochlorococcus* strains, which is known to have the smallest genome of any free-living photosynthetic cell (Biller et al., 2014).

**Figure 4.**
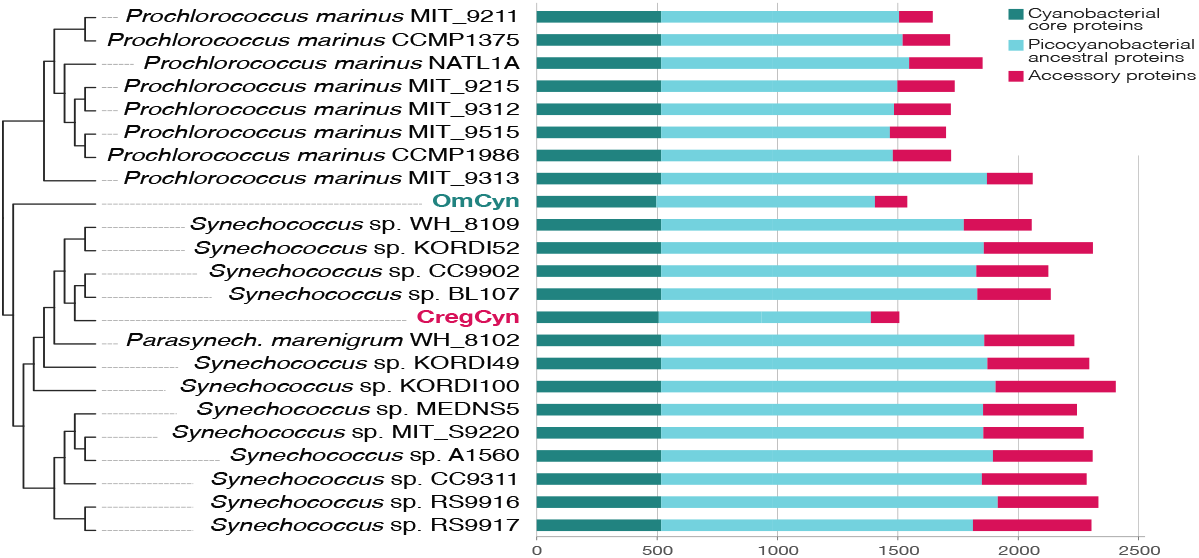
Nonredundant proteome size comparisons of cyanobacterial genomes. The comparison is based on the number of orthologous protein groups encoded in each genome. Each bar represents the proteome size without functional redundancy. The cladogram on the left shows the phylogenetic relationships in Figure 2.

Since CregCyn and OmCyn are likely to have been acquired independently, the reduction in protein repertoire is also likely to have occurred independently. In contrast to OmCyn, for which there is no phylogenetically related lineage with a known genome sequence, CregCyn is clearly closely related to the free-living strains of the clade IV for which genome sequencing has been completed, and the reduction in protein repertoire is apparent (i.e., *Synechococcus* sp. CC9902 and BL107, with protein repertoire sizes of 2125 and 2135, respectively; Figure 4). To investigate repertoire reduction in different protein categories, we classified each protein group into ‘core proteins’, ‘picocyanobacterial ancestral proteins,’ and ‘accessory proteins’ based on their degree of conservation in the phylogeny (see Materials and Methods) and repertoire reduction in each category was examined. The results showed that the protein repertoire reduction of CregCyn is also similar to that of OmCyn, and that almost all of the reduction occurred in the ‘picocyanobacterial ancestral proteins’ and ‘accessory proteins’ categories (Figure 4).

To further investigate the qualitative differences between the CregCyn and OmCyn genome reductions, we compared the proteins encoded by these two symbiont genomes with the predicted ancestral picocyanobacterial proteome to understand which proteins were lost in each symbiont lineage (Supplementary figure S4). The picocyanobacterial ancestral proteome inferred in this study consists of 2,369 protein groups, of which 1,157 could be functionally annotated using the KEGG orthology (Kanehisa et al., 2016). The number of orthologous protein groups identified in the CregCyn and OmCyn genomes was 1,388 and 1,405, respectively. Both correspond to approximately 59% of the ancestral proteome, again suggesting that a similar degree of proteome reduction from the ancestral proteome occurred in both symbiont genomes. The similarity in proteome reduction between the two lineages was confirmed not only in the number of remaining proteins, but also in their functional breakdown; 1,259 protein genes, that are representing approximately 90% of the ancestral protein genes remaining in the CregCyn and OmCyn genomes (90.7% for CregCyn, 89.6% for OmCyn), were shared between these independently evolved dinoflagellate symbionts (Supplementary figure S4a). This trend was also observed for functionally annotated proteins; functionally annotated ancestral proteins common to CregCyn and OmCyn (850 protein genes) accounted for 93.6% and 94.1% of all functionally annotated proteins found in their respective genomes (Supplementary figure S4a). Investigation of the number of lost proteins by functional category showed no marked bias (Supplementary figure S4b). The similar genome reduction trends in CregCyn and OmCyn, two cyanobacterial symbionts with different origins, can be attributed to their shared lifestyle; These symbionts reside in unique extracellular chambers of the dinoflagellate hosts rather than intracellularly. Notably, the ‘transporters’ category exhibited the largest number of lost proteins in both symbionts (Supplementary figure S4b), potentially due to their adaptation to the stable and less variable environment within the symbiotic chamber, in contrast to the external environment. Neither symbiont contained genes related to ‘Bacterial motility proteins’ (Supplementary figure S4b), supporting the idea that they remain stationary within the chambers throughout their life cycle. On the other hand, as shown in previous studies, compared to the extent of genome reduction in other endosymbiotic cyanobacteria or in cyanobacteria-derived organelles other than ordinary plastids (e.g., the chromatophore of *Paulinella* spp. (Lhee et al., 2017; Lhee et al., 2019; Nowack et al., 2008)), the genome reduction in CregCyn and OmCyn is very mild, and these symbiotic cyanobacteria appear to maintain their metabolic independence. This fact also makes the genome evolution of the symbiotic cyanobacteria of Dinophysiales dinoflagellates unique.

### Evolutionary difference between CregCyn and OmCyn

The molecular phylogenetic analysis of CregCyn and OmCyn not only suggests that the two cyanobacteria have different origins, but also provides insight into the timing of the establishment of symbiosis with their respective hosts. Since CregCyn is shown to be a sister lineage to the marine *Synechococcus* subcluster 5.1 clade IV, it is likely that the ancestral cyanobacterium of CregCyn established the symbiotic relationship with dinoflagellates after most of the diversity of marine *Synechococcus* lineages seen today had emerged. OmCyn, on the other hand, is a lineage that branches from the base of subcluster 5.1, and to our knowledge, there are no data supporting the presence of free-living strains in this lineage. Given that the OmCyn lineage has not been found in previous comprehensive diversity analyses of marine picocyanobacteria targeting free-living cells, it could be expected that all species of the lineage currently represented by OmCyn have symbiotic relationships with dinoflagellates and that the establishment of this symbiotic relationship predates the diversification of the marine *Synechococcus* subcluster 5.1, which is estimated to have occurred approximately 240 million years ago (Sánchez-Baracaldo et al., 2019).

### Genomic islands in the CregCyn genome

Our comparative analysis of protein repertoires shows that despite their independent origins, both CregCyn and OmCyn genomes have undergone similar reductive evolution in parallel. However, further close examinations of genome traits revealed notable distinctions between the two symbiotic cyanobacterial genomes. We assessed the tetranucleotide diversity spectrum of each genomic region within a genome, which is considered one of the common metrics for picocyanobacterial genome analysis (Dufresne et al., 2008; Scanlan et al., 2009). Heterogeneity in tetranucleotide frequency across genomic regions has been observed in the marine picocyanobacteria and serves as an indicator of ‘genomic islands’ that correspond to the clusters of putative exogenous genes and contribute to adapting to habitats with different environmental parameters (Juhas et al., 2009; Scanlan et al., 2009). Our analysis unveiled distinct tetranucleotide frequency profiles in certain regions of the CregCyn genome, similar to its free-living relatives (*Synechococcus* sp. CC9902 and BL107, both from the clade IV; Figure 5). Conversely, the region-wise tetranucleotide frequency profiles of the OmCyn genome showed relatively low diversity (Figure 5). If the CregCyn genome subregions with biased tetranucleotide frequencies are genomic islands, these are expected to contain proteins not conserved in closely related cyanobacteria, implying horizontal gene transfer (HGT). Considering this idea, we attempted to score the possibility of the HGT origin of each protein-coding gene found in the CregCyn genome and assess their relationship with tetranucleotide frequencies for each genomic subregion. First, we performed sequence homology searches for each of all CregCyn proteins against databases containing protein sequences from a broad range of bacterial diversity, fully covering cyanobacterial lineages. The organisms that appeared as hits in the search results were surveyed, and the frequencies of hits for each organism were calculated; Proteins from cyanobacteria closely related to CregCyn were the most frequent, as expected. We then used this frequency data to assign scores to the hit organisms for each protein in its homology search. The score will be high if organisms that appear as hits in a protein’s homology search are predominantly those with a high hit frequency in the genome-wide analysis and low if these organisms are rare in the genome-wide homology search. As a result, several genomic regions exhibited notably low frequency scores of hit organisms, which can be considered as a proxy for the possibility of foreign origin, and some of these regions also contained peaks of biased tetranucleotide frequencies (Figure 6). We regarded these overlapped regions as ‘genomic islands’ in the CregCyn genome in this study (refer to Supplementary table S1 for a list of protein-coding genes located on the genomic islands).

**Figure 5.**
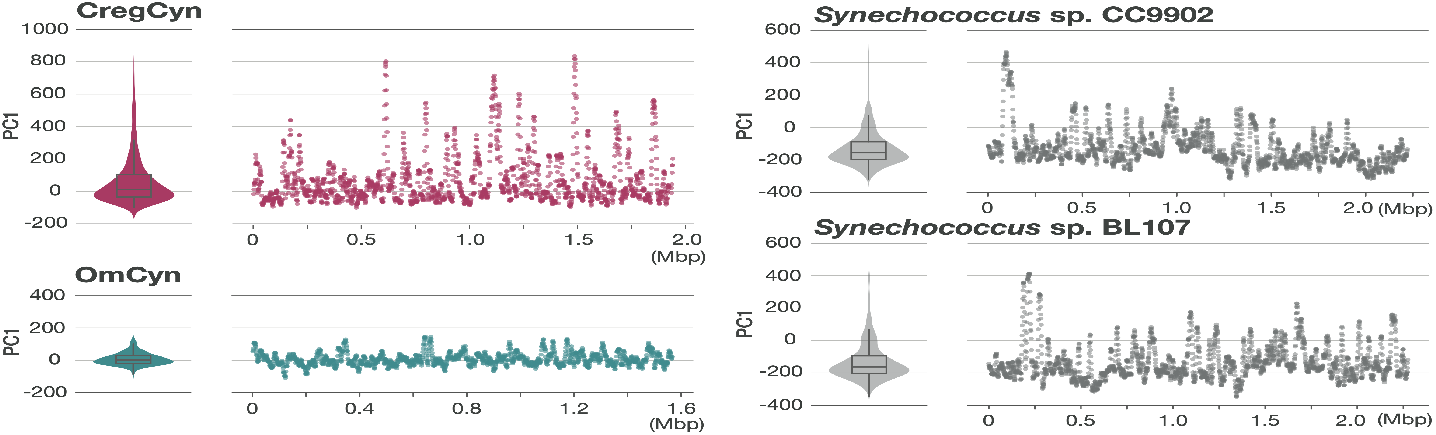
Genome-wide diversity of tetranucleotide frequency of cyanobacterial genomes. The tetranucleotide frequencies for each genomic subregion were obtained using a sliding window approach with a width of 10 Kbp and an overlap of 1 Kbp. The frequency values for a genome were analyzed by using principal component analysis and the values in the first principal component (PC1) were used as summary values for the tetranucleotide frequency. Violin plots with box plots show the overall diversity of tetranucleotide frequencies of the genomes. Scatter plots show PC1 values for each subregion in a genome.

**Figure 6.**
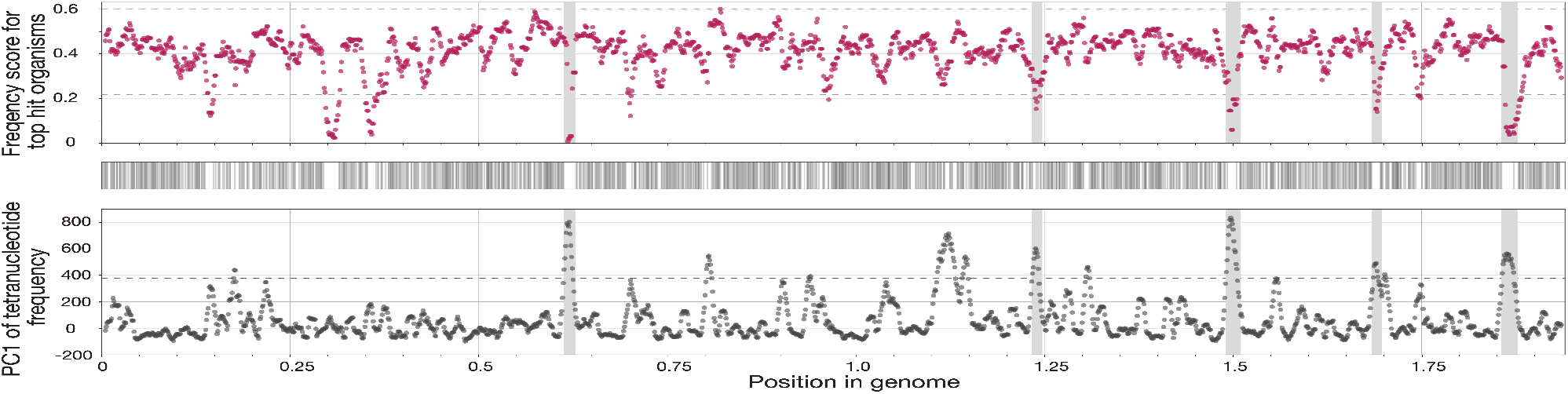
Genomic islands detected in the CregCyn genome. Scatter plot on top: average frequency score of top-hit organisms for proteins located within each genomic subregion. These subregions were defined using the same windows as those employed in the bottom plot. Plot on middle: presence of *Synechococcus* sp. CC9902, BL107 and *Parasynechococcus* marenigrum in hit organisms for each homology search; these organisms were the most common among the hit organisms in the overall homology search for all of the CregCyn proteins. Scatter plot on bottom: the values on the first principal component (PC1) from the principal component analysis, illustrating the tetranucleotide frequencies for each genomic subregion; this data was also presented in Figure 5. Horizontal dotted lines in both the top and bottom scatter plots indicate values corresponding to the mean plus or minus two times the standard deviation of the data. Genomic islands (shaded areas) were defined as subregions where values exceeding this threshold were observed in both plots.

As discussed above, OmCyn may have continued its symbiotic relationship with dinoflagellates prior to the current diversity of the marine *Synechococcus* subcluster 5.1 being formed. Assuming that the lineage has survived only within the dinoflagellate chamber during this time, it is likely that the OmCyn lineage has had very limited contact with organisms in the external environment, consistent with the fact that the low diversity of tetranucleotides found and the lack of detectable genomic islands in this genome. On the other hand, it is noteworthy that genomic islands are present in the CregCyn genome despite its reductive nature. However, the timing of the origin of these genomic islands, which is potentially important in the context of the evolution of the symbiotic relationship, is unknown. One of the possible explanations for the presence of genomic islands in CregCyn is that it has spent less time since the establishment of the symbiotic relationship with the dinoflagellate host than that of OmCyn. It is possible that the current genomic islands in the CregCyn genome already existed in the free-living ancestor of CregCyn, and no new genomic islands have been formed since then. However, not enough time has passed for the genomic islands to become undetectable through acclimation. On the other hand, given that the coexistence of multiple cyanobacteria and other bacteria in a dinoflagellate chamber has been reported (Foster et al., 2006a), we cannot rule out the possibility of HGT within the chamber. In any case, the currently available data do not allow us to speculate when these genomic islands were formed, whether before or after CregCyn established a symbiotic relationship with dinoflagellates and whether genes associated with the genomic islands contributed to the adaptation to the symbiotic lifestyle. Future comparisons with the symbiotic cyanobacterial genomes of other Citharistes species may provide further insight into this question.

## Conclusions

In this study, we first succeeded in revealing the complete genome sequence of a cyanobacterial symbiont of the pelagic dinoflagellate *C. regius*. Through genome analysis, the evolutionary and metabolic characteristics of the symbiont were revealed, as well as similarities with the genome of OmCyn, another cyanobacterial symbiont of dinoflagellate with different origins. However, the specific role of CregCyn in this symbiotic relationship, including the mechanism of metabolite exchange between CregCyn and *C. regius*, remains unclear. Addressing these biological questions will require continued efforts involving challenges for laboratory cultivation of this symbiotic consortium, close cell observation, as well as comprehensive sequence analysis. Another important outcome of this study is the presentation of another complete genome of marine picocyanobacteria with unique genomic features, such as a reductive protein repertoire. Based on several indications from previous research, it is expected that the diversity of cyanobacteria symbiotically associated with Dinophysiales dinoflagellate is even greater (Foster et al., 2006b). Further genomic analysis of these cyanobacteria will reveal their genomic and genetic diversity.

## Materials and Methods

### Sampling of *Citharistes regius* cells and genome amplification of symbionts

The cells of *Citharistes regius* used in this study were found in a seawater sample collected with a plankton net (mesh size: 25 μm) off Shimoda, Shizuoka Prefecture, Japan (N34°29.222’, E139°06.200’) on November 15, 2021. Three *C. regius* cells found in a sample (cell IDs: A11, A12, and M16; Supplementary figure S2) were picked up with a microcapillary, washed more than three times by transferring droplets of sterilized seawater, and finally placed in sterilized fresh water for the subsequent genome amplification step. Each of the three *C. regius* cells harboring cyanobacterial symbionts was subjected to independent genome amplification using REPLI-g Single Cell Kit (QIAGEN), and the amplified DNA samples were processed with S1 nuclease (TaKaRa) to reduce branching junctions.

### Genome sequencing, assembling, and annotation

Illumina sequencing libraries for three amplified DNA samples were prepared using Nextera TruSeq DNA PCR-Free Low Throughput Library Prep Kit (Illumina). Subsequently, the libraries were analyzed on an Illumina NovaSeq 6000, yielding 33 to 43 million paired-end reads per library, each measuring 150 bp in length. The quality control of the Illumina short reads was performed using fastp (version 0.12.4; Chen et al., 2018) with default settings. To obtain long-reads, one of the amplified genomic samples (M16) was further subjected to library construction using the Rapid Sequencing Kit (Oxford Nanopore Technologies), and the library was analyzed on a MinION platform (Oxford Nanopore Technologies) with a MinION Flow Cell (R9.4.1). The MinION sequencing generated 70 thousand nanopore reads, totaling approximately 162 Mbp. The resulting reads had an average length of 2.3 Kbp, a maximum length of 55 Kbp, and an N50 of 4.5 Kbp.

The genome of M16 cell was assembled *de novo* in a hybrid manner, incorporating both the Illumina short reads and the nanopore long reads. This assembly was carried out using Unicycler (version 0.4.8; Wick et al., 2017) with the ‘–no_pilon’ option. Additional genome assembling for A11 and A12 cells utilizing their Illumina short reads were also conducted using Unicycler. Preliminary genome annotations for all three assemblies were performed using DFAST (version 1.2.15; Tanizawa et al., 2018). For the predicted proteins obtained through the preliminary annotation of the M16 cell assembly, a sequence homology search against the refseq_proteins database (O’Leary et al., 2016) was conducted, leading to the identification of four contigs encoding protein genes with high similarity to those of cyanobacteria. The same analysis was conducted for the assemblies using short reads of M11 and M12 cells, but no additional cyanobacterial contigs were identified beyond those observed in the M16 assembly. An analysis of the assembly graphs for the M16 assembly using Bandage (version 0.8.1; Wick et al., 2015) revealed that three of the four contigs form a single graph, while the remaining contig of 17 Kbp in size forms a circular DNA molecule. Sequence gaps between the three cyanobacterial contigs were closed by PCR, revealing a circular main chromosome of 1.94 Mbp in size. The annotation of the main chromosome and a small circular DNA molecule (plasmid) resulting from the above procedure was initially performed by DFAST in an automated fashion, followed by manual curation to obtain the final annotation. KEGG Orthology ID (KO ID) assignment to each protein was performed by using the KEGG Automatic Annotation Server (Moriya et al., 2007) with the BBH (bi-directional best hit) method.

### Ortholog detection and protein repertoire analysis

Orthologous relationships of the predicted proteins of CregCyn with proteins from other cyanobacterial lineages were estimated using OrthoFinder (version 2.5.3; Emms & Kelly, 2019). This analysis utilized proteomes from nine cyanobacterial symbionts, including CregCyn and OmCyn, as well as proteins observed in a phylogenetically diverse range of cyanobacterial genomes, totaling 252 complete genomes (see Supplementary table S2 for the genome list). The cyanobacterial proteins were clustered based on their similarity by OrthoFinder and grouped into orthogroups, each containing orthologous proteins. The size of the non-redundant protein repertoire for each genome was defined as the total number of orthogroups assigned to proteins within their proteomes, plus the number of proteins not assigned to any orthogroup (species-specific proteins).

Orthogroups conserved in all free-living cyanobacterial genomes analyzed in this study were designated as ‘core proteins’. In addition, ‘picocyanobacterial ancestral proteins’ were defined as orthogroups found in both the *Prochlorococcus* clade and the *Synechococcus* subcluster 5.1. Proteins that did not fall into either of these categories were identified as “accessory proteins”.

### Phylogenetic analysis

#### 16S rRNA gene phylogeny

For the phylogenetic analysis of CregCyn based on the 16S rDNA sequence, homologous sequences of cyanobacteria were collected from public databases. This dataset consisted mainly of the sequences from the *Synechococcus*/*Prochlorococcus*/*Cyanobium* clade, of which *Synechococcus elongatus* occupies a basal position, and it also included partial cyanobacterial 16S rRNA gene sequences (286-379 bp) reported by Foster et al. (2006b), which were found in reverse transcription products of whole *Citharistes* sp. cells from natural environments. The 16S rRNA gene sequences of CregCyn and other cyanobacteria were aligned using the L-INS-i method of MAFFT (version 7.49; Katoh & Standley, 2013) and trimmed using trimAl (version 1.4; Capella-Gutiérrez et al., 2009) with ‘–gt 0.8’ option. Phylogenetic analysis was conducted on the trimmed alignment using IQ-TREE (version 2.1.2; Minh et al., 2020) with the TIM3+F+R3 model, which was estimated as the best-fit model using the model test tool in IQ-TREE (Kalyaanamoorthy et al., 2017). Support values for nodes were evaluated using 100 nonparametric bootstrap replicates under the same substitution model.

#### Phylogenomic analysis using multiple protein sequences

To perform phylogenetic analysis using multiple protein sequences, protein sequences of 143 single-copy genes conserved in 99% of picocyanobacterial genomes were extracted based on the result of the OrthoFinder analysis. The extracted protein sequences were aligned by orthogroups using the L-INS-i method of MAFFT (version 7.49; Katoh & Standley, 2013), and all sites with gaps in the alignments were removed using trimAl (version 1.4; Capella-Gutiérrez et al., 2009). These 143 protein datasets were combined into a final dataset consisting of 23 sequences and 35,518 sites. This dataset is available upon request. The final dataset was subjected to the maximum likelihood phylogenetic analysis using IQ-TREE (version 2.1.2; Minh et al., 2020) with the LG+C20+F+Γ4 substitution model. Bootstrap values were obtained from a nonparametric bootstrap analysis with 100 replicates. The LG+C20+F+Γ4 PMSF model (Wang et al., 2018) was used for the bootstrap analysis, with the maximum likelihood tree topology as the guide tree.

### Relative abundance estimation using metagenomic data

Metagenomic data from the *Tara* Oceans Project (Pesant et al., 2015; Sunagawa et al., 2015) were used to estimate the relative abundance of each cyanobacterial genome in natural environments based on size fraction. 593 metagenomic samples obtained from 68 *Tara* Oceans sampling sites were downloaded from the Sequence Read Archive (SRA) of the National Center for Biotechnology Information (see Supplementary table S3 for accession IDs of SRA data used). Five million reads from each metagenome were mapped to each cyanobacterial genome using BLASTN (version 2.12.0+; Zhang et al., 2000) of the BLAST+ package, and metagenomic reads that showed*≥* 99% identity to a single genomic sequence spanning a minimum of 90 bases were treated as corresponding reads for the specific genome. In this process, the reads mapped to coding genomic regions encoding rRNA and tRNA were excluded from the analysis, as these regions have high sequence homology across cyanobacterial lineages. The reads corresponding to each genome found in each metagenome were summed for each size fraction, i.e., 0.8-5 μm, 5-20 μm, 20-180 μm, and 180-2,000 μm. The relative abundance of each genome was determined by calculating the ratio of the summed read counts to the total number of metagenomic reads subjected to mapping for each size fraction.

### Genomic island detection

#### Tetranucleotide frequency analysis

The comparison of tetranucleotide frequencies between subregions of a genome was performed as follows. The occurrence of all possible tetranucleotides was assessed for each genomic window obtained using a sliding window approach with a width of 10 Kbp and an overlap of 1 Kbp. The counts for each tetranucleotide across genomic windows were combined to generate a 2-dimensional dataset comprising vectors of tetranucleotide counts for each window. The tetranucleotide dataset was centered by subtracting its mean value. Principal component analysis (PCA) was then performed on the centered dataset. The first principal component from the PCA results was used to assess diversity and bias in tetranucleotide frequencies across genomic windows. The main chromosome of the genome of each organism was used for this analysis. If the genome sequence was represented by several scaffolds, as is the case for OmCyn, the longest scaffold was used in the analysis.

#### Estimation of the origin of each CregCyn protein

To assess the frequency scores of top hit organisms in the homology search for each CregCyn protein, we first conducted a DIAMOND BLASTP homology search (version 2.0.9; Buchfink et al., 2021) for all CregCyn proteins against the refseq protein database (O’Leary et al., 2016) as well as the OmCyn total protein dataset (Nakayama et al., 2019). From the DIAMOND-BLASTP results for the all CregCyn proteins, we determined hit frequencies for each organism in the database by examining the top five hits for each protein. Hits to sequences of “unidentified *Synechococcus*” were excluded from the count in this step. These hit frequency values of organisms were then normalized using the min-max normalization approach, transforming them to a range from 0 (minimum value) to 1 (maximum value). For each CregCyn protein, the top five hit organisms in the DIAMOND-BLASTP results were evaluated by referencing the normalized frequency values of hit organisms. The mean of the frequency value for these five hits was defined as the frequency score for the respective CregCyn protein. Finally, to compare the frequency scores with the tetranucleotide frequency trends in each genomic subregion, the mean frequency scores for protein-coding genes within each subregion were calculated using the same sliding window approach. Protein-coding genes that partially overlap boundary regions of a window are also incorporated into the calculation as belonging to that window. In addition, for *Synechococcus* sp. CC9902, *Synechococcus* sp. BL107, and *Parasynechococcus marenigrum*, which were the most frequent hit organisms in the overall homology search for all CregCyn proteins, we checked whether these three strains appeared in the top five hits of the homology search for each protein, and the results are plotted in Figure 6 (middle plot).

To evaluate the variability in tetranucleotide frequency trends and hit organism frequency scores within each genomic region, regions with values exceeding the threshold of the mean plus or minus two times the standard deviation of the individual data were treated as peaks of interest. Subregions that exceeded the threshold in both tetranucleotide frequency data and hit organism frequency scores were designated as genomic islands in this study.

## Acknowledgement

This work was supported by JSPS KAKENHI Grant Numbers JP16H06280, 20H03305, 18KK0203, 23H02535, and BPI05044. We thank the *Tara* Oceans consortium and sponsors who supported the *Tara* Oceans Expedition for making the data accessible. Computations in this study were partially performed on the NIG supercomputer at ROIS National Institute of Genetics. The manuscript file uploaded to bioR*χ*iv was generated using the LaTeX template adapted by Stephen Royle (https://github.com/quantixed) available at https://github.com/quantixed/manuscript-templates.

## Data availability

The annotated genome data of CregCyn has been deposited in DDBJ (the DNA DataBank of Japan)/GenBank/EMBL (European Molecular Biology Laboratory) under the accession numbers AP029048 and AP029049.

