## Supplementary figure S1 for "Convergent reductive evolution of cyanobacteria in symbiosis with Dinophysiales dinoflagellates"

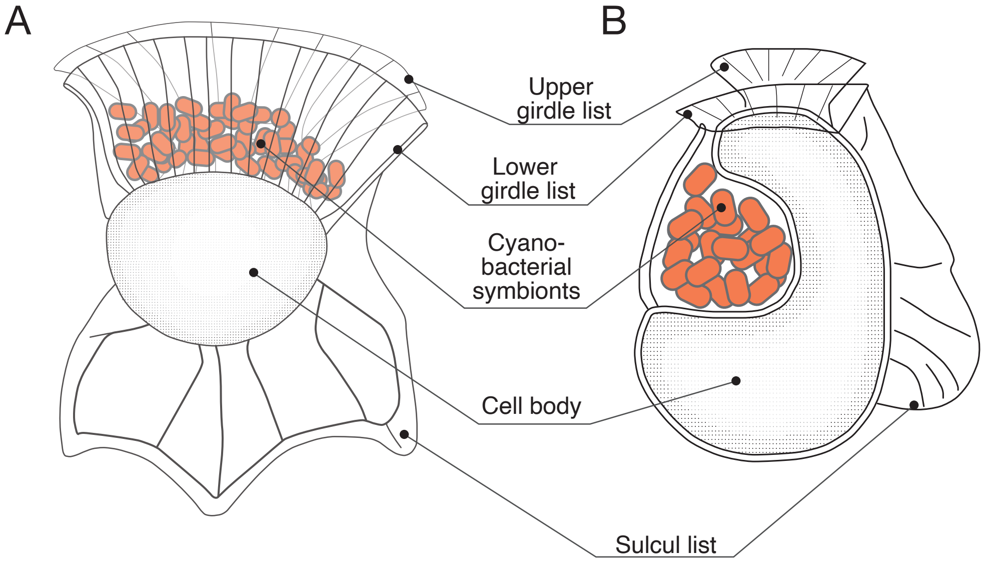


**Figure S1.** Schematic drawing of the morphology of *Ornithocercus magnificus* (A) and *Citharistes regius* (B).
