## Supplementary figure S2 for "Convergent reductive evolution of cyanobacteria in symbiosis with Dinophysiales dinoflagellates"

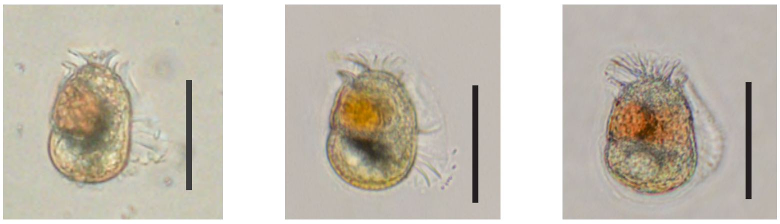

A11 A12 M16

**Figure S2.** Micrographs of *Citharistes regius* individuals used for whole genome amplification in this study. Scale bars = 40μm.
