## Supplementary figure S3 for "Convergent reductive evolution of cyanobacteria in symbiosis with Dinophysiales dinoflagellates"

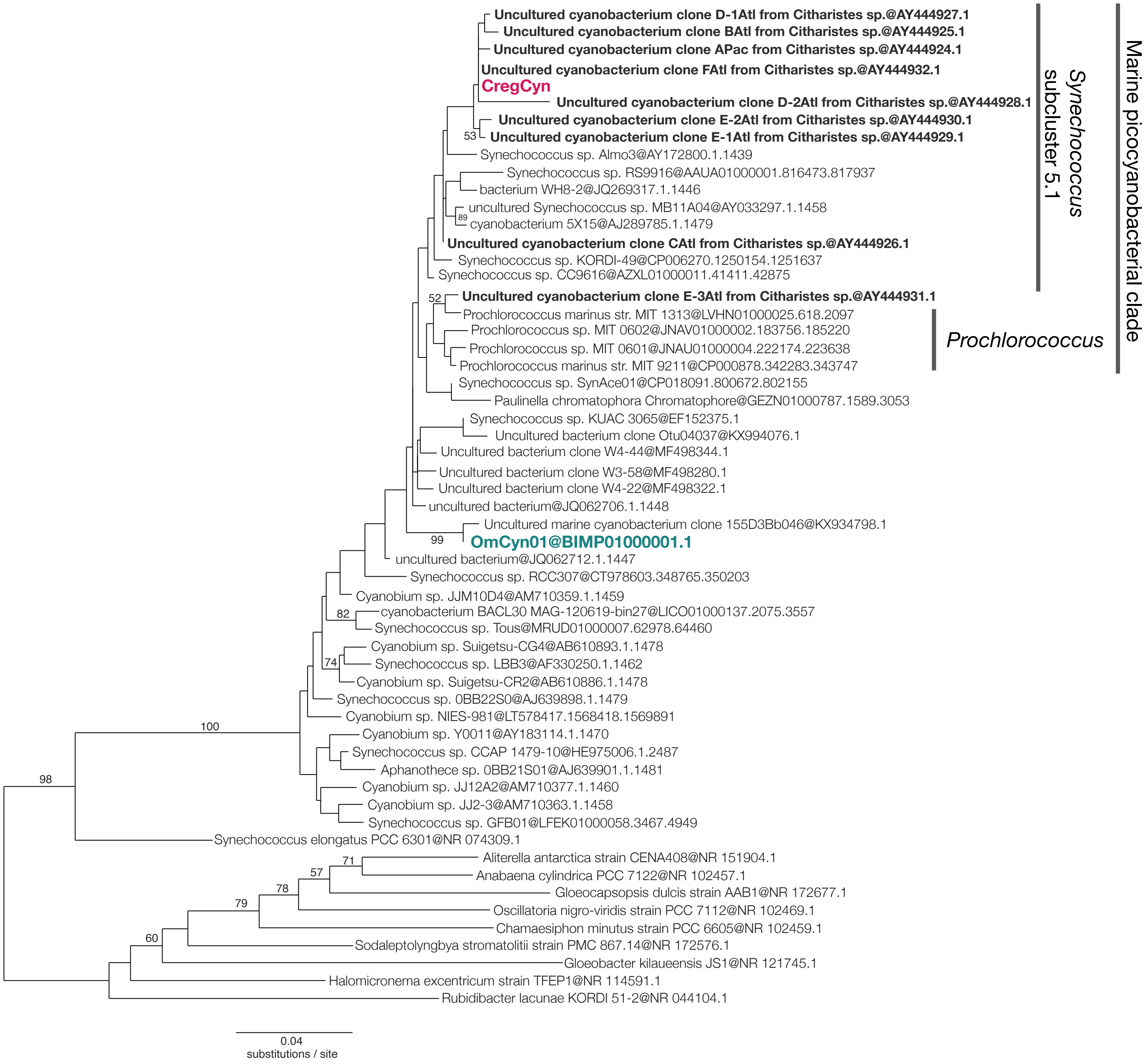


**Figure S3.** Maximum likelihood phylogenetic tree inferred from 16S rRNA gene sequences. The *Synechococcus/Prochlorococcus* clade is displayed as ingroup. CregCyn and OmCyn sequences are shown in magenta and green, respectively. Other sequence labels in bold are partial sequences obtained from cells of *Citharistes* spp. that are reported by Foster et al. 2006. The numbers shown for each branch are bootstrap support values. Only bootstrap values of 50 or higher are shown. The scale bar represents the estimated number of nucleotide substitutions per site.
