## Supplementary figure S4 for "Convergent reductive evolution of cyanobacteria in symbiosis with Dinophysiales dinoflagellates"

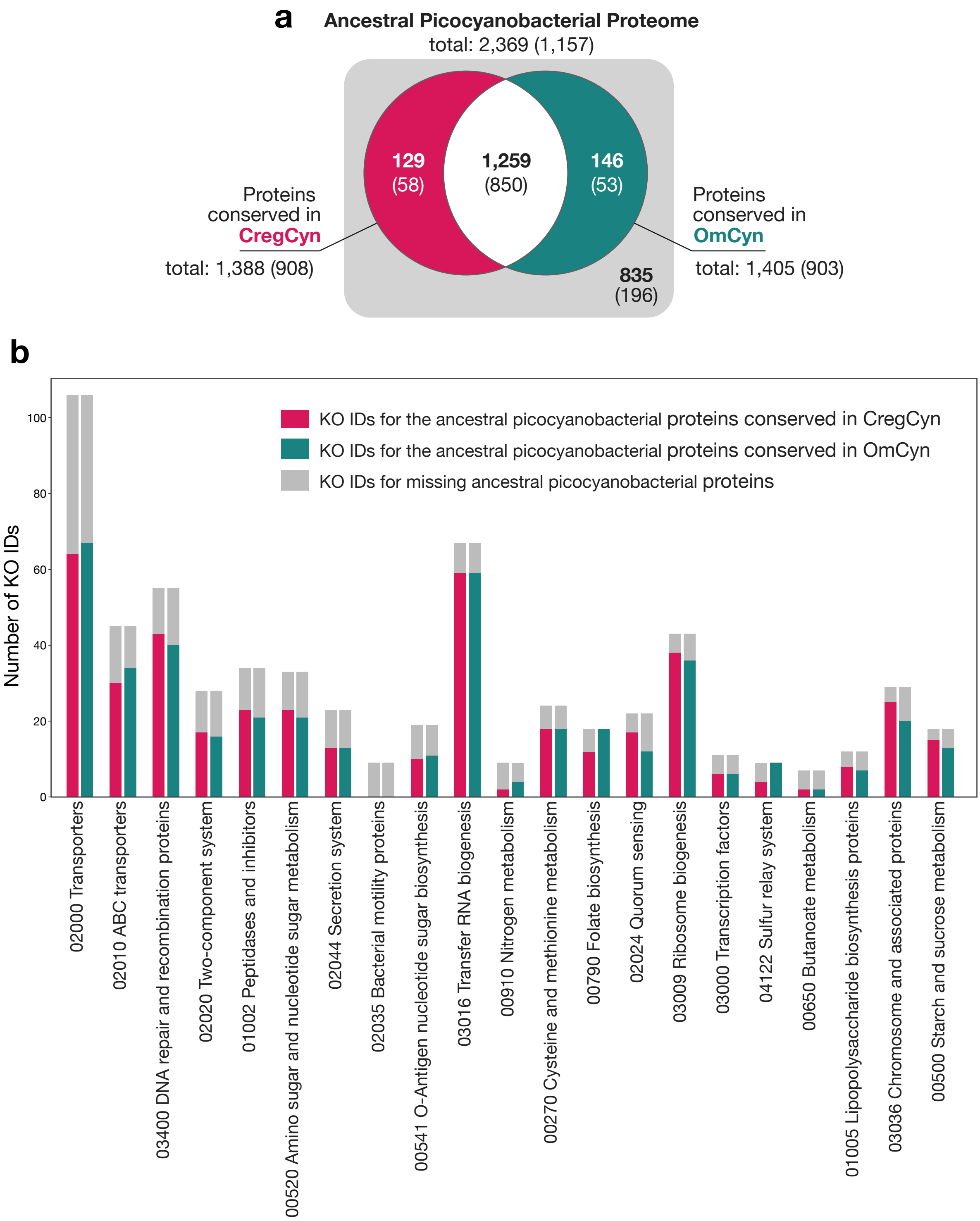


**Figure S4.** Ancestral picocyanobacterial proteins remaining in the CregCyn and OmCyn genomes. (a) Venn diagram of total ancestral picocyanobacterial proteins found in the CregCyn and OmCyn genomes. The numbers in the diagram indicate the number of proteins. Numbers in parentheses indicate the number of proteins that have KEGG Orthology IDs assigned. (b) Breakdown by KEGG functional categories.
